# Changes in cerebral glucose metabolism at rest following hemicontusive spinal cord injury in mice: A whole-brain autoradiographic study

**DOI:** 10.1101/2025.10.01.679887

**Authors:** Zhuo Wang, Camelia Danilov, Dilan Setiya, Daniel P. Holschneider

**Author notes:** Correspondence to: Daniel P. Holschneider, MD, Department of Psychiatry and the Behavioral Sciences, University of Southern California, 1975 Zonal Avenue, KAM 400, MC9037, Los Angeles, CA 90089-9037, USA. These authors contributed equally (Zhuo Wang, Camelia Danilov).

## Abstract

Spinal cord injury (SCI) disrupts brain-spinal cord communications and results in profound brain reorganization. Here, we apply high-resolution, voxel-based, whole-brain metabolic mapping using the [^14^C]-2-deoxyglucose autoradiographic method in mice to assess functional brain reorganization in a subacute stage (1 week after SCI). Right moderate contusive injury at the cervical 5 level (C5) was confirmed by glial fibrillary acidic protein (GFAP) immunohistochemical staining. SCI compared to sham-lesioned animals showed significant motor deficits (grip strength, rotarod) alongside decreases in glucose uptake in sensorimotor regions of the cortex, basal ganglia, and thalamus, which receive monosynaptic afferents (the ventral posterolateral thalamic nucleus, VPL) or multi-synaptic afferents from the spinal cord (the primary somatosensory and motor cortices, caudate putamen). In contrast, regions in the limbic system (the amygdala, accumbens nucleus, lateral septum, and hippocampus) and in the cerebellum demonstrated increases in glucose uptake in SCI animals. Most of these effects were noted bilaterally, suggesting functional reorganization involving higher order neural circuits bilaterally. The current findings underscore the broadness of brain reorganization in the subacute stage following incomplete SCI. Functional whole-brain metabolic mapping provides a roadmap for future targeted studies examining neuroplastic mechanisms in search of new therapeutic strategies.

## INTRODUCTION

Following spinal cord injury (SCI), the central nervous system undergoes significant reorganization not only in the spinal but also in the supraspinal circuits (Solstrand Dahlberg *et al*., 2018). After an incomplete SCI, intraspinal projection neurons can generate new “detour circuits” that transmit information onto the spinal cord from above to below the injury (Murray *et al*., 2010; Takeoka *et al*., 2014) (Filli & Schwab, 2015). Corticospinal tract fibers can regenerate, sprout, or reorganize to recover functional circuits alongside remaining neurons in the spinal cord (Oudega & Perez, 2012). In addition, following SCI, the control of movement is not limited to the corticospinal tract fibers, but may also recruit other descending pathways such as the reticulospinal, rubrospinal, and tectospinal tracts (Calderone *et al*., 2024). Supraspinal reorganization is proposed to be the result of neuroplastic changes with complex spatial and temporal patterns following SCI, including acute changes in response to the damage of efferent fibers and deafferentation, as well as long-term changes involving retrograde neuronal degeneration, afferent fiber remodeling, and adaptation at the circuit level (see review (Calderone *et al*., 2024)). Contributing plasticity includes the growth of new axonal branches (Jain *et al*., 2000; Fouad *et al*., 2001; Weidner *et al*., 2001; Vavrek *et al*., 2006), synaptic remodeling (Li *et al*., 2004; Kim *et al*., 2006), homeostatic synaptic plasticity, and changes in neuronal membrane characteristics such as sustained inward currents (Harvey *et al*., 2006) among others.

Clinical and preclinical neuroimaging studies on SCI have advanced our understanding of structural (using computer tomographic imaging CT, magnetic resonance imaging MRI) and functional changes in the brain (using resting-state and task-related functional MRI). These studies demonstrate widespread changes in structure, regional activation, somatosensory and motor topographic maps, and inter-regional functional connectivity while details of these findings may vary depending on the type, severity, and stage of SCI (reviewed by Wang et al. 2019, Nardone et al. 2013) (Nardone *et al*., 2013; Wang *et al*., 2019). Fewer studies have examined changes in brain metabolism, which is an important marker for function. While clinical studies using [^18^F]-fluoro-deoxyglucose (FDG) positron emission tomography (PET) typically examine patients in the chronic stage, often months to years after SCI, studies in rodent models of SCI are relatively few. Jaiswal et al. using [^18^F]-FDG PET in rats have demonstrated a reduction in glucose uptake in the cerebellum 90 days after bilateral thoracic SCI (Jaiswal *et al*., 2022). Horner et al (1995) applied cerebral autoradiography to a rat model of bilateral thoracic SCI and analyzed a limited number of regions-of-interest 3 months after injury. They noted a reduction in glucose uptake in the ventral posterolateral thalamic nucleus (VPL), cerebellar lobules 7 and 8, and the nucleus gracilis (Horner & Stokes, 1995). To our best knowledge, a whole-brain analysis of changes in glucose metabolism during the subacute phase of SCI has not been conducted. In the current study, we apply high-resolution, voxel-based, whole-brain autoradiographic mapping of glucose uptake 1 week after hemicontusive cervical SCI, at a time of functional brain reorganization before changes have become chronic.

## MATERIALS & METHODS

Female C57/BL mice were purchased from Charles River (Hollister, CA, USA) at 8 - 10 weeks of age. Animals were housed on a 12-hour light/12-hour dark cycle (lights on from 7:00 am to 7:00 pm) in groups of four with direct bedding and with free access to water and standard rodent chow. All procedure and handling techniques were in strict accordance with the National Institutes of Health guidelines for the care and use of laboratory animals and approved by the Institutional Animal Care and Use Committee of the University of Southern California, with accreditation by the Association for Assessment and Accreditation of Laboratory Animal Care (AAALAC) International. Studies were initiated after 1 week of acclimatization to the vivarium.

### Spinal cord surgical procedures

The SCI model used in this study was a unilateral contusion injury model at the 5th cervical level (C5) as previously described (Aguilar & Steward, 2010), with the change that the tip impactor was positioned on the right side of the cord, creating primarily unilateral deficits to the right forelimb (Danilov *et al*., 2020). Mice were deeply anesthetized through intraperitoneal injection of a cocktail of ketamine (100 mg/kg) and xylazine (10 mg/kg), and a dorsal laminectomy was performed on C5 (*n* = 6). The contusion injury was created using the Infinite Horizon device platform (Precision Systems and Instrumentation, LLC, Fairfax Station, VA, USA). A moderate injury was achieved using an impactor (diameter = 1.3 mm) and an impact force of 80 kDyn.Control animals (Sham) were anesthetized and received skin incision only (*n* = 7). Mice received a single dose of Ethiqa XR (3.25 mg/kg, s.c., MWI Animal Health, Concora Inc., Boise, ID, USA) prior to the surgery for analgesia, and a single dose of enrofloxacin (2.5 mg/kg, s.c., MWI Animal Health) post-surgery for prophylactic treatment against urinary tract infection. For 3 days post-surgery, mice received an injection of Ringer’s lactate solution daily (1 ml/20 g, s.c.) for hydration. Mice were housed individually following surgery.

### Behavioral evaluation

SCI-induced motor deficits were assessed using the grip strength and accelerating rotarod test. In the grip strength test, a stainless-steel wire cage lid (wire diameter = 0.2 cm, inter-wire distance = 0.8 cm) was covered with duct tape around the edges to prevent the mouse from climbing around the rim, leaving a 6 cm x 10 cm window with exposed wires. On post-surgery day 6, the animal was tested in three trials with a 30-min inter-trail interval. In each trial, the mouse was placed on top of the lid where wires were exposed. The lid was slightly tilted to force the animal to grip the wires and then quickly turned upside down. The lid was held 40 cm from the floor, which was covered with 1-cm of bedding. The time-to-fall was recorded with a stopwatch manually with a 180-s cut-off time. The mean was calculated after removing the shortest time-to-fall.

In the rotarod test, the animal was trained, and baseline measured over 3 days before the surgery. The animal was placed on the spindle (diameter = 7.3 cm) of the rotarod (Economex Rotarod, Columbus Instruments, Columbus, OH, USA) for 30 s followed by 5 min walking at 5 rpm (115 cm/min) for two days. On the 3^rd^ day, baseline was measured in 3 trials with a 30-min inter-trial interval. In each trial, the time-to-fall was recorded manually with a starting speed of 5 rpm, and an acceleration of 0.1 rpm/s (2.3 cm/min/s). The cut-off time was 300 s. The mean was calculated after removing the shortest time-to-fall. The animal was then tested on post-surgery day 6 and the mean was normalized to the baseline.

Data were presented as mean ± standard error of mean for each group and analyzed using GraphPad Prism 10 (GraphPad Software, LLC., Boston, MA, USA). If the data passed the Shapiro-Wilk normality test, two-tailed Student’s *t*-test was used to evaluate between-group differences. Otherwise, two-tailed Mann-Whitney test was used. *P* < 0.05 was considered statistically significant.

### Measurement of cerebral glucose uptake at rest (Holschneider *et al*., 2019)

On day 4 – 6 post-surgery, the mice were habituated to the experiment room with dim lighting for 15 min daily. On day 6, the mice were fasted overnight with free access to water. On day 7 post-surgery, each mouse was administered i.p. [^14^C]-2-deoxyglucose (2DG, 0.3 µCi/g bodyweight in 0.53 ml saline, American Radiolabeled Chemicals, Inc., St. Louis, MO, USA), followed by a 45-min radiotracer uptake period in a resting state in home cage. Thereafter, mice were deeply anesthetized with 5% isoflurane and euthanized by cervical dislocation.

### Autoradiography

Brains were extracted and flash frozen in methylbutane over dry ice (about - 55 ºC). Brains were then serially cryosectioned into coronal slices (20-μm thick, 140-μm inter-slice distance) at -18 ºC in a cryostat (Mikron HM550, Thermo Fisher Scientific, Waltham, MA, USA). Slices were heat-dried on glass slides and exposed to UltraCruz autoradiography film (Santa Cruz Biotechnology, Inc., Dallas, TX, USA) alongside [^14^C] radioactivity standards (Amersham Biosciences, Piscataway, NJ, USA). Autoradiograms of brain slices were digitized on an 8-bit grey scale using a voltage-stabilized light box (Northern Light R95 Precision Illuminator, Imaging Research Inc., St. Catharines, Ontario, Canada) and a Retiga 4000R charge-coupled device monochrome camera (QImaging of Teledyne Photometrics, Tucson, AZ, USA). Regional cerebral glucose uptake (rCGU) was measured according to the method of Sokoloff (Sokoloff *et al*., 1977) with modification (Melzer *et al*., 1985; Jordan *et al*., 2005; Vyazovskiy *et al*., 2008). Spatial resolution of 2-DG autoradiographic imaging is ∼50 μm (Stumpf & Solomon, 1995).

### Image Processing (Nguyen *et al*., 2004)

3-D brains were reconstructed from 70 consecutive autoradiograms (approximate bregma levels: + 1.54 mm to – 6.72 mm, voxel size: 40 μm x 140 μm x 40 μm) using TurboReg (Thevenaz *et al*., 1998) and a nonwarping geometric model (rotations, translations and nearest-neighbor interpolation). A brain template was created using our prior methods (Thevenaz *et al*., 1998), with background and ventricular spaces thresholded based on optical density. Spatial normalization consisted of a 12-parameter affine transformation followed by a nonlinear spatial normalization using 3-D discrete cosine transforms. Final normalized brains were smoothed with a Gaussian kernel (full-width at half-maximum = 240 μm × 420 μm × 240 μm) to improve the signal-to-noise ratio. To account for global differences in absolute amount of [^14^C], voxel intensities of each brain were proportionally scaled.

### Analysis of regional cerebral glucose uptake (rCGU) and metabolic connectivity (Holschneider *et al*., 2019)

Statistical parametric mapping (Friston *et al*., 1995; Nguyen *et al*., 2004) (SPM5, Wellcome Centre for Neuroimaging, University College London, London, UK) was used for data analysis. Unbiased, voxel-by-voxel Student’s *t*-tests between the SCI and Sham group were performed across the whole brain to access changes in rCGU following SCI. Threshold for statistical significance was set at *P* < 0.05 at the voxel level with an extent threshold of 200 contiguous voxels, which reflects a balanced approach to control Type I and Type II errors. Regions were identified by a mouse brain atlas (Franklin & Paxinos, 2007).

We applied seed-ROI (region of interest) correlation analysis (Holschneider *et al*., 2014) to assess metabolic connectivity of the ventral posterolateral thalamic nucleus (VPL), a part of the sensory thalamus that receives monosynaptic input from the contralateral spinal cord. A circular, structural ROI was hand drawn in MRIcro (version 1.40, http://cnl.web.arizona.edu/mricro.htm) in the left VPL at bregma – 1.58 mm over the template brain according the mouse brain atlas. Mean optical density of the seed ROI was extracted for each animal using the MarsBaR toolbox for SPM (version 0.42, http://marsbar.sourceforge.net/). Correlation analysis was performed in SPM for each group using the seed values as a covariate. Threshold for significance was set at *P* <0.05 at the voxel level and an extent threshold of 200 contiguous voxels. Regions showing significant correlations (positive or negative) in rCGU with the seed ROI are considered functionally connected with the seed.

### Immunohistochemistry

Immunohistochemistry was performed as previously described (Danilov & Steward, 2015). Mice were euthanized with an overdose of ketamine and xylazine (160 and 20 mg/kg bodyweight, respectively) and perfused transcardially with 4% paraformaldehyde in 0.1 M sodium phosphate buffer (Na_2_HPO_4_, pH 7.4). Spinal cords were dissected and post-fixed in 4% paraformaldehyde overnight, then immersed in 30% sucrose for cryoprotection overnight, frozen in TissueTek OCT (VWR International, Radnor, PA, USA) and stored at − 20°C until sectioning with a cryostat. The tissue block extending from ∼ 4 mm above to 4 mm below the lesion and containing the injury site was sectioned at 20 µm intervals in the horizontal plane. The sections were stained with specific antibodies for glial fibrillary acidic protein (GFAP Dako, Cat# Z0334). After blocking in 5% normal goat serum in PBS solution, sections were incubated in primary antibody (GFAP 1:1000) at 4 ^0^C overnight. The following day, sections were washed in PBS and incubated with the fluorescent secondary antibody (anti-rabbit Alexa Fluor 488, 1:250) for 2 hrs at RT. After twice rinsing in PBS, sections were mounted on gelatin subbed slides. Coverslips were applied with VECTASHIELD, Antifade Mounting Medium with DAPI (Cat # H-1200, Vector Laboratories, Burlingame, CA, USA). Images were obtained on a BZ-9000 (BIOREVO) fluorescence microscope (Keyence Corporation of America, Itasca, IL, USA).

## RESULTS

Confirmation of unilateral lesion of the cervical spinal cord is shown in **Fig. 1**. Immunohistochemical staining confirmed tissue loss in the right hemicord with superimposed image showing increases in GFAP immunostaining in the perilesional area. SCI compared to Sham animals demonstrated statistically significant deficits in the grip strength test (time-to-fall: SCI, 10.4 ± 8.0 s, *n*= 6; Sham, 171.8 ± 8.2 s, *n* = 7. *P* = 0.001, Mann-Whitney test) and the accelerating rotarod test (time-to-fall: SCI, 17.9 ± 7.0 %; Sham, 102.1 ± 5.7 % of baseline. *P* < 0.0001, Student’s *t* test), suggesting SCI-induced impairment in strength, coordination and balance (**Fig. 2**).

**Fig. 1.**
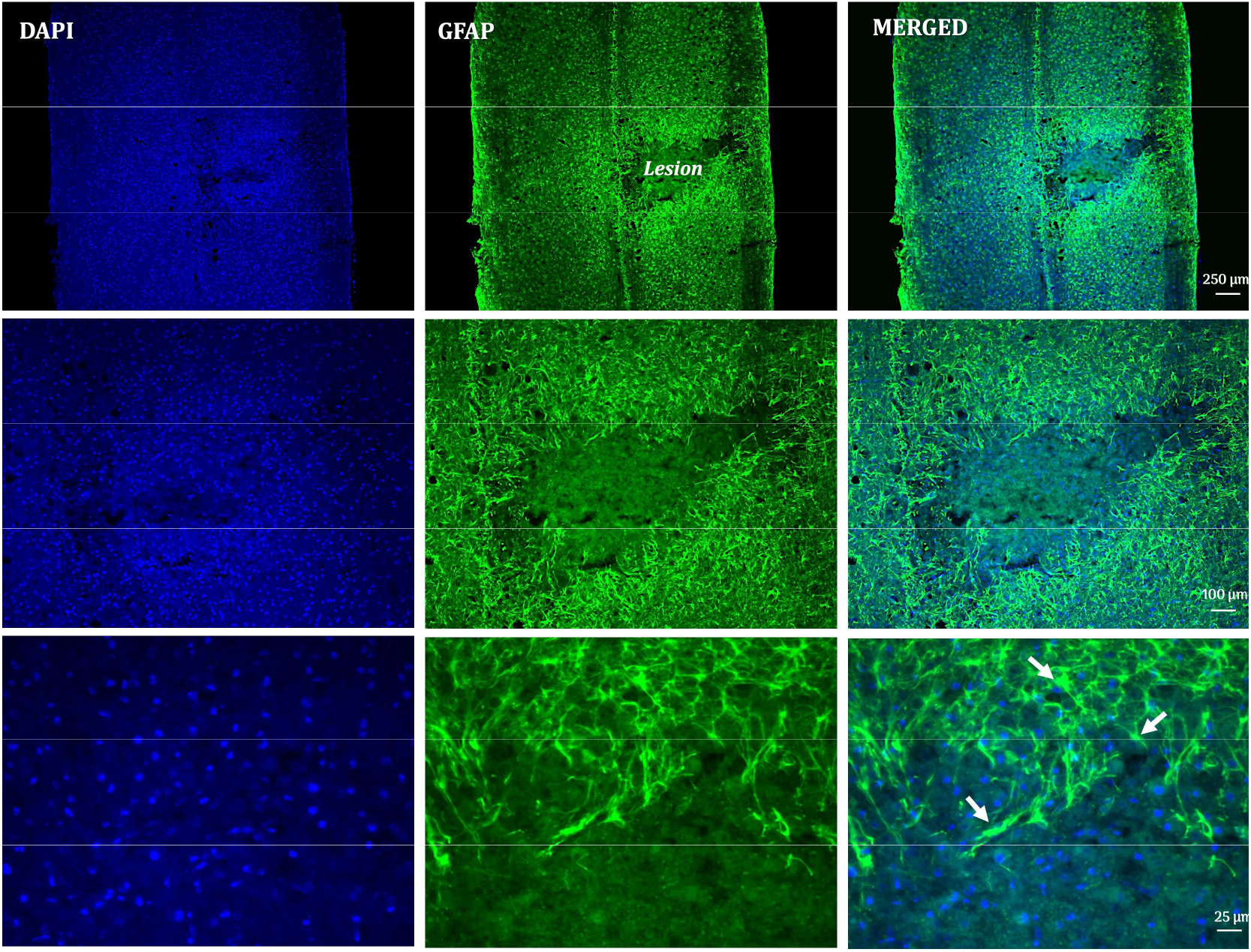
Representative images of right unilateral lesion after an 80 kDyn contusive cervical spinal cord injury in mice. Arrows indicate increased expression of glial fibrillary acidic protein (GFAP) in reactive astrocytes in response to injury.

**Fig. 2.**
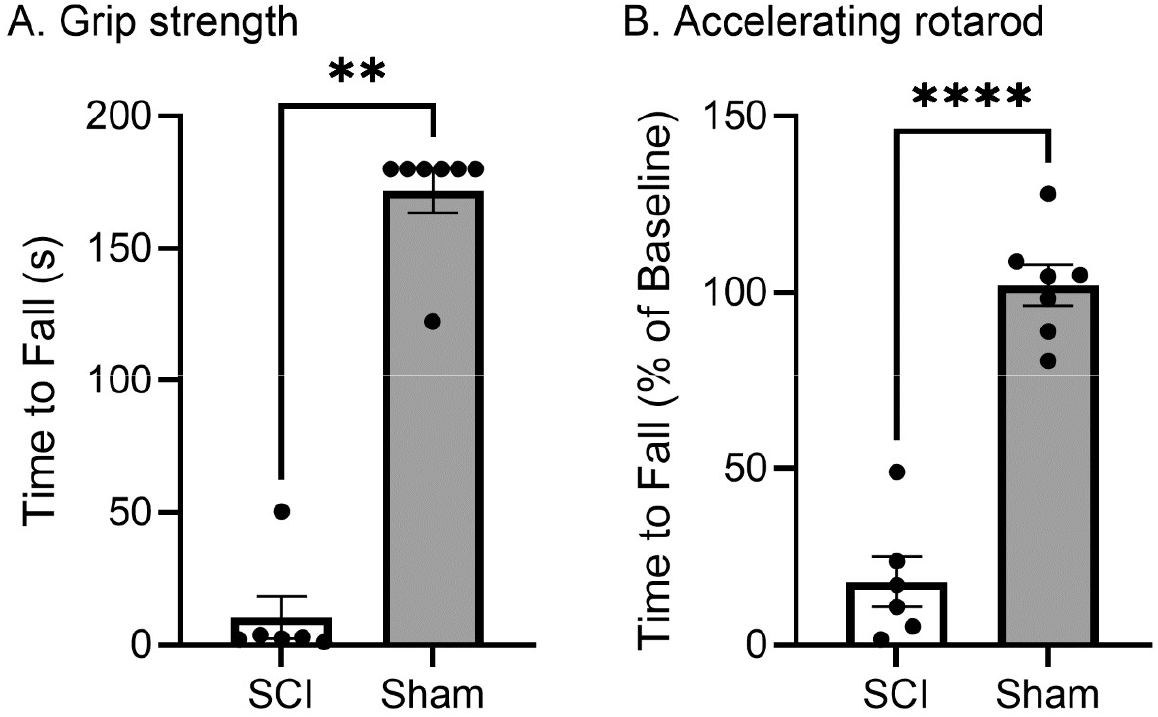
Motor deficits after spinal cord injury (SCI). The spinal cord injury (SCI) mice showed statistically significant motor deficits compared to the Sham mice in the grip strength test (A, **: *P* = 0.001, Mann-Whitney test) and the accelerating rotarod test (B, ****: *P* < 0.0001, Student’s *t* test, *n* = 6 for SCI, 7 for Sham group).

Statistically significant changes in rCGU were noted broadly in the SCI mice compared to Sham at rest 1 week following cervical hemicontusive SCI (**Table 1, Fig. 3)**. It is noteworthy that most of the SCI-related changes in rCGU were bilateral. SCI-related decreases in rCGU were seen mostly in the anterior part of the brain (rostral to ∼ bregma - 5.0 mm) including the primary and secondary somatosensory (S1, S2), primary and secondary motor (M1, M2), piriform (Pir), insular (I) cortices, basal ganglia (caudate putament CPu, globus pallidus GP, left substantia nigra SN, left subthalamic n. STh), thalamic nuclei (anterodorsal n. AD, habenula n. Hb, laterodorsal n. LD, mediodorsal n. MD, paraventricular n. PV, posterior n. Po, ventral posterolateral n. VPL; ventral posteromedial n. VPM), and left mesencephalic reticular formation (mRt).

**Table 1:** Regions showing statistically significant differences in regional cerebral glucose uptake in mice at rest 1 week following cervical hemicontusive spinal cord injury compared to sham mice.

|  | Contralesional | Ipsilesional |
| --- | --- | --- |
| **CORTEX** |  |  |
| Auditory (Au) | + | ++ |
| Entorhinal, dorsolateral (DLEnt) | + | ++ |
| Insular (anterior to posterior, I) | - | - |
| Motor, primary (M1) | -- | -- |
| Motor, secondary, anterior (M2) | - | - |
| Motor, secondary, posterior (M2) | + | + |
| Parietal association (PtA) | ++ | + |
| Piriform (Pir) | - | - |
| Somatosensory, primary, anterior (S1) | - | - |
| barrel field (S1BF) | - | - |
| jaw (S1J) | - | - |
| forelimb (S1FL) | -- | - |
| upper lip region (S1ULp) | - | - |
| Somatosensory, secondary (S2) | -- | - |
| Visual, primary/secondary (V1/V2) | + |  |
| **SUBCORTEX** |  |  |
| Accumbens n., core (AcbC) |  | ++ |
| shell (AcbSh) | + | ++ |
| Amygdala, basolateral, basomedial n. (BL, BM) | + | + |
| medial n., anterior part (MeA) | + | + |
| posteromedial cortical n. (PMCo) |  | + |
| Caudate putamen, ventral (vCPu) | -- | -- |
| dorsal (dCPu) | - | - |
| central (cCPu) | -- | -- |
| Cerebellum, crus 1 of the ansiform lobule (Crus1) | + | + |
| interposed cerebellar n. (Int) | + |  |
| lateral cerebellar n. (Lat) | + | + |
| lobules 4/5 of the cerebellar vermis (4/5Cb) | + | + |
| middle cerebellar peduncle (mcp) | + | + |
| Globus pallidus (GP) | - | - |
| Hippocampus, CA1 | + | + |
| CA2 | + | + |
| CA3 | ++ | + |
| dentate gyrus (DG, posterior) | ++ | + |
| Hypothalamus, posterior (PH) |  | + midline |
| Lateral septum (LS) | + | + |
| Parabrachial pigmented n. of the ventral tegmental area (PBP) | - |  |
| Parasubiculum (PaS) |  | + |
| Pons, reticular n. (PnC, PnO) | + | + |
| Reticular formation, mesencephalic (mRt) | - |  |
| Reticular n., intermediate (IRt) |  | + |
| Substantia nigra (SN) | -- | - |
| Thalamic nuclei |  |  |
| anterodorsal (AD) | - | - |
| anteromedial (AM) |  | + |
| habenula (Hb) | - | - |
| laterodorsal (LD) | -- | -- |
| mediodorsal (MD) | - | - |
| paraventricular (PV) |  | - midline |
| posterior (Po) | -- | - |
| reticular (Rt) |  | ++ |
| subthalamic n. (STh) | - |  |
| ventral posterior lateral (VPL) | -- |  |
| ventral posterior medial (VPM) | -- | - |
| Tegmental n., laterodorsal (LDTg) | + | + |
| Trigeminal motor n. (5N) | + | + |
| Ventral pallidum (VP) | + | + |
| Vestibular n. (Ve) | + | + |
| Zona incerta (ZI) | - |  |

Significant increases or decreases in regional cerebral glucose uptake are noted with '+' and '-', respectively, for the contra- and ipsilesional hemispheres. Significance is shown at the voxel level for $\geq 200$ contiguous voxels at + / - $P < 0.05$ and ++ / -- $P < 0.01$ . Abbreviations are taken from the Franklin and Paxinos mouse brain atlas (Franklin & Paxinos, 2007).

**Fig. 3.**
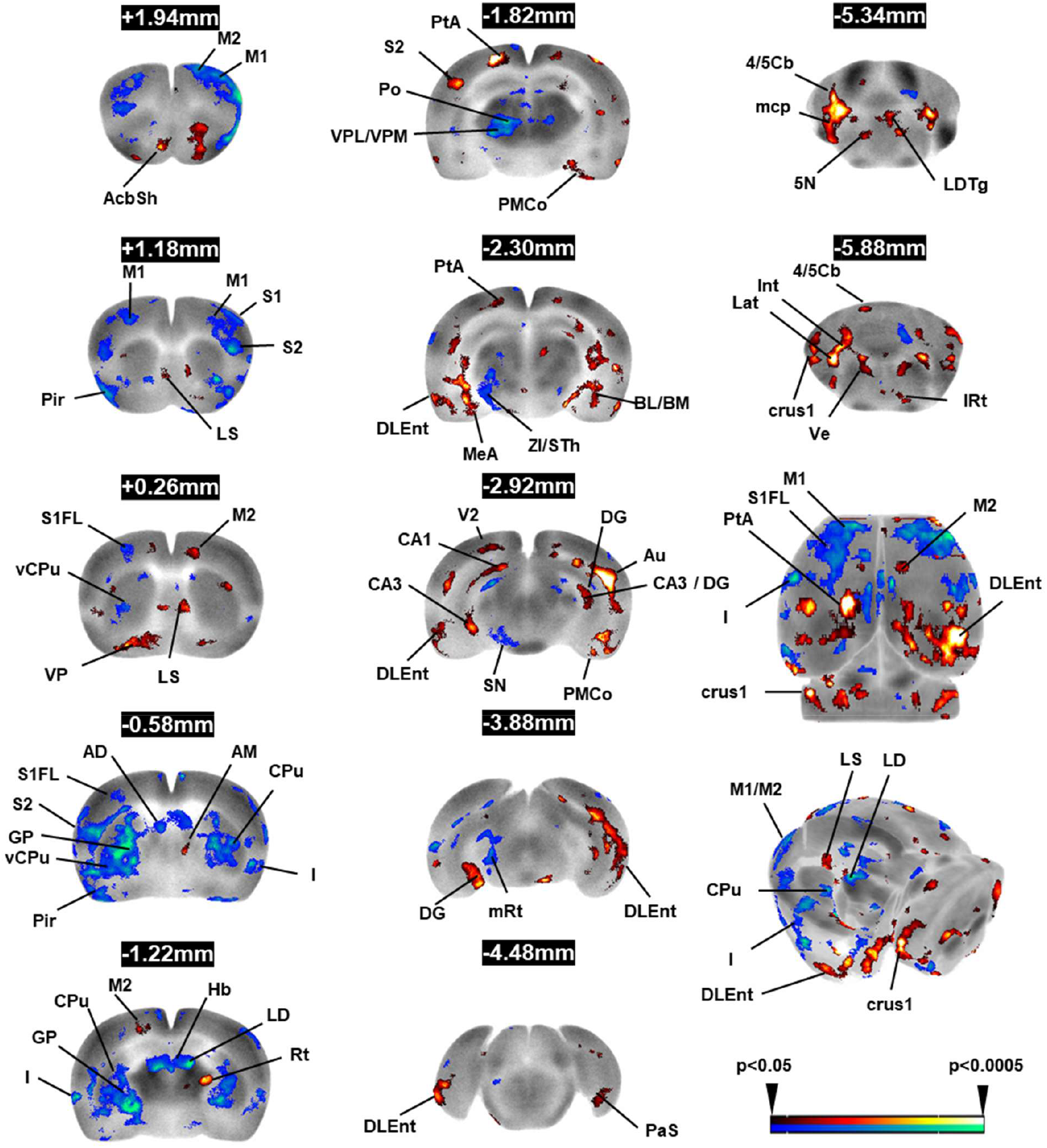
Effects of right, hemicontusive cervical spinal cord injury (SCI) on regional cerebral glucose uptake (rCGU) in the resting state 1 week following injury. Significant changes in rCGU in SCI mice compared to sham are shown. Representative coronal sections of the template brain (anterior-posterior level relative to the bregma shown in black textbox) depict the ipsilesional (right) and contralesional (left) hemispheres. The color-coded overlays are statistical parametric maps showing significant changes in rCGU (red/blue: increases/decreases, *P* < 0.05, >200 contiguous voxels, *n* = 6-7 per group). The bottom two images of the 3^rd^ column show significant changes near the surface of the 3-D rendered template brain. *Abbreviations* Acb, accumbens nucleus AD, anterodorsal thalamic n. Au, auditory cortex BL, basolateral amygdaloid n. BM, basomedial amygdaloid n. CA1/CA2/CA3, hippocampus CPu, striatum Crus1, crus of the ansiform lobule DG, dentate gyrus DLEnt, dorsolat.entorhinal cortex GP, external globus pallidus Hb, habenular n. I, insular cortex LD, lateral dorsal thalamic n. LDTg, laterodorsal tegmental n. LS, lateral septum M1/M2, primary/secondary motor cortex mcp, middle cerebellar peduncle MeA, medial amygdaloid n. PaS, parasubiculum Pir, piriform cortex PMCo, cortical amygdaloid n. Po, posterior n. thalamic n. PtA, parietal association cortex Rt, reticular thalamic n. S1/S2, primary/secondary somatosensory cortex SN, substantia nigra STh, subthalamic n. V2, secondary visual cortex Ve, vestibular n. VL, ventrolateral thalamic n. VM, ventromedial thalamic n. VP, ventral pallidum VPL, ventroposterior lateral thalamic n. VPM, ventroposterior medial thalamic n. ZI, zona incerta 4/5 Cb, lobules 4 & 5 of the vermis 5N, trigeminal motor n.

In contrast, SCI-related increases in rCGU were noted in many limbic regions including the amygdala (basolateral n. BL, basomedial n. BM, medial n. anterior part MeA, posteromedial cortical n. PMCo), accumbens nucleus (shell AcbSh, right core AcbC), lateral septum (LS), posterior hypothalamus (PH), and hippocampus (CA1, CA2, CA3, dentate gyrus DG, parasubiculum PaS, dorsolateral entorhinal cortex DLEnt). Increases in rCGU were also seen in the cortex (posterior M2, S2, auditory Au, parietal association PtA, left primary/secondary visual V1/V2), cerebellum (lobules 4 and 5 of the vermis Cb4/5, interposed n. Int, lateral n. Lat, middle cerebellar peduncle mcp, crus 1 of the ansiform lobule Crus1), ventral pallidum (VP), thalamus (right anteromedial n. AM, right reticular n. Rt), and brainstem areas (parabrachial pigmented n. of the ventral tegmental area PBP, pontine reticular n. oral/caudal part PnO/PnC, right intermediate reticular n. IRt, laterodorsal tegmental n. LDTg, motor trigeminal n. 5N, vestibular n. Ve).

The VPL is a major relay center for ascending input from the spinal cord and showed decreases in rCGU following SCI primarily on the contralesional side. We further applied seed correlation analysis to assess SCI-related changes in metabolic connectivity of the VPL. A seed representing the contralesional (left) VPL was selected at bregma -1.82 mm. **Fig. 4**. compares metabolic connectivity of the VPL seed in the SCI and the Sham group. Compared to the Sham group, the SCI group showed greatly reduced positive metabolic connectivity with the motor (M1, M2), primary somatosensory (including forelimb S1FL, hindlimb S1HL, shoulder S1Sh, and barrel field S1BF areas), parietal association, and visual (V1/V2) cortices, as well as with the cerebellum (lobule 3 of the vermis 3Cb, simple lobule Sim, Crus1). The SCI group also showed negative correlation with the retrosplenial cortex (RS) and 3Cb, which were absent in the Sham group. There was also attenuation of intrathalamic connectivity in SCI compared to Sham mice (data not shown).

**Fig. 4.**
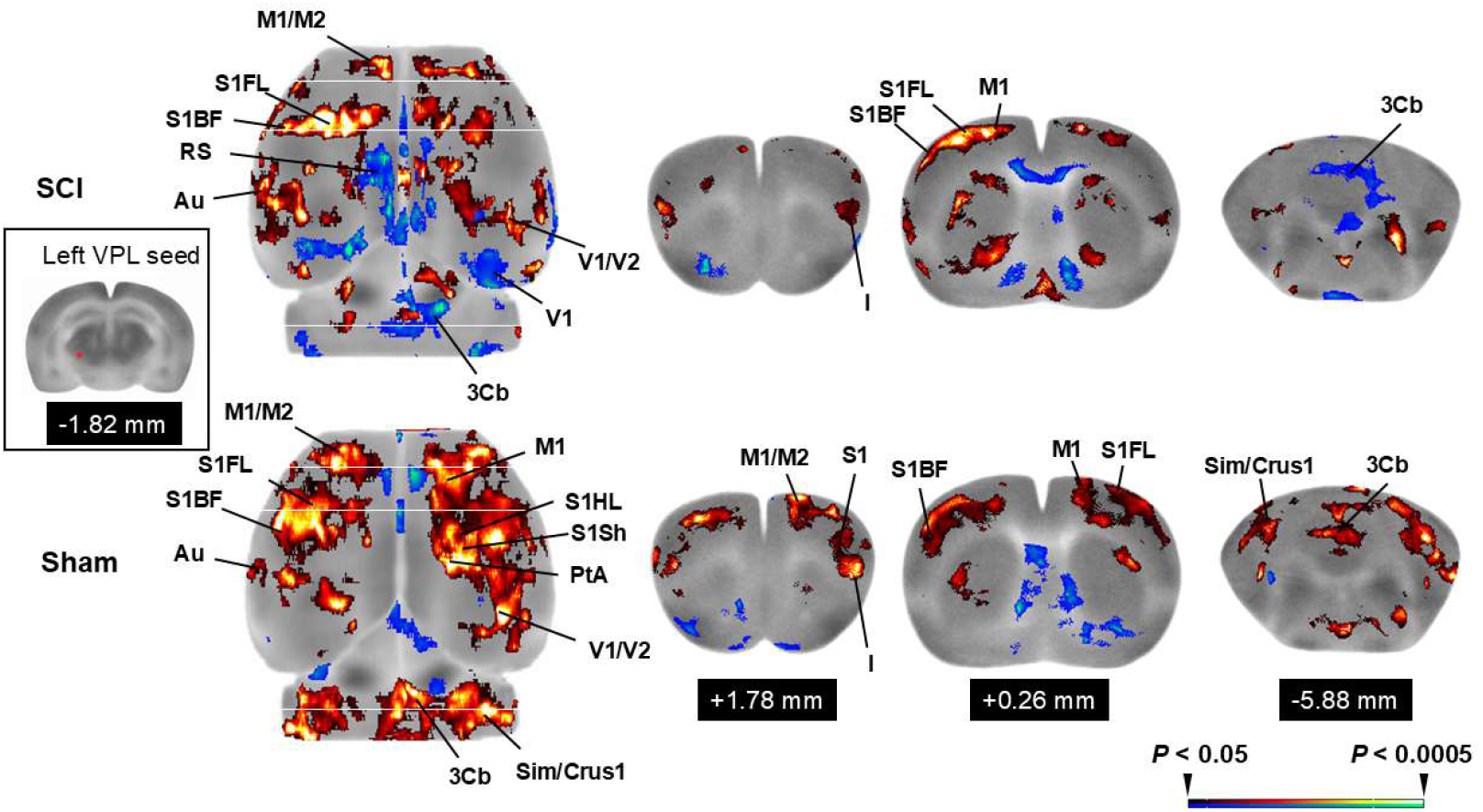
Changes in metabolic connectivity of the contralesional ventral posterolateral thalamic nucleus (VPL) to the cortex and cerebellum following hemicontusive cervical spinal cord injury (SCI). A seed representing the contralesional (left) VPL was selected at bregma -1.82 mm (inset on the left). The transverse view of the 3D rendered template brain is shown alongside three coronal sections the bregma levels of which are shown in the black textboxes and marked on the transverse figure with white horizontal lines (top panel: SCI; lower panel: Sham). Color-coded overlays show brain regions that have statistically significant correlations in regional cerebral glucose uptake with the VPL seed (red/blue: positive/negative correlation. *P* < 0.05, >200 contiguous voxels, *n* = 6-7 per group). For the 3D rendered images, visible depth of overlay was set to 1 mm to reveal cortical connectivity. Compared to the Sham group, the SCI group showed greatly reduced metabolic connectivity with motor and somatosensory cortices, a loss of positive correlation with the cerebellum, and newly emerged negative correlation with the retrosplenial and posterior visual cortices. Abbreviations: 3Cb (lobule 3 of the cerebellar vermis), Au (auditory cortex), Crus1 (crus 1 of the ansiform lobule), I (insular cortex), M1/M2 (primary/secondary motor cortex), PtA (parietal association cortex), RS (retrosplenial cortex), S1 (primary somatosensory cortex), S1BF (S1 barrel field), S1FL (S1 forelimb), S1HL (S1 hindlimb), S1Sh (S1 shoulder), Sim (simple lobule), V1/V2 (primary/secondary visual cortex).

## DISCUSSION

Our results indicate extensive changes in regional cerebral glucose uptake in animals 1 week after hemicontusive cervical SCI compared to normal controls. Changes were associated with significant motor deficits on the grip strength test and accelerating rotarod test as previously reported (Danilov *et al*., 2020). Cerebral metabolic changes were regionally specific. In general (with exceptions as noted below), relative glucose uptake was decreased in forebrain motor sensory regions, as well as in regions of the basal ganglia and sensory thalamus. In contrast, regions in the limbic system and in the cerebellum demonstrated a relative increase in glucose uptake.

A notable finding following the hemicontusive cervical SCI was the extent of bilateral changes in glucose metabolism. For example, hypometabolism was seen in the forelimb area of S1 in both the contralesional and ipsilesional hemisphere. Hemicontusive SCI at the cervical level has been shown to cause damage to both the gray and white matter (including the dorsal column, and to some extent the lateral and ventral funiculi). While a great majority of descending fibers, e.g. the corticospinal tract, originate in the contralateral hemisphere and cross the midline before reaching the spinal cord, different ascending fibers passing through the cervical segments target brain regions in *both* hemispheres. Fibers in the dorsal column cross the midline in the medulla and target the contralateral hemisphere. Other ascending fibers, e.g. spinothalamic tract, coming from the lower segments cross the midline below the cervical level and target the ipsilateral hemisphere. Damaged spinal neurons in the cervical segments can cause deafferentation of regions in the contralateral hemisphere. Therefore, hemicontusive cervical SCI can affect both hemisphere depending on the severity of lesion. Such profound reorganizations of the ipsilesional sensory-motor cortex after unilateral spinal cord injury has been previously reported in rats (Ghosh *et al*., 2009). Likewise, functional MRI (fMRI) in rodents has previously shown that while unilateral paw stimulation in normal animals results largely in activation of contralateral somatosensory cortex, in spinal cord-injured animals, there is a much more extensive activation, which includes contralateral but also ipsilateral structures such as cortex, thalamus, hippocampus, and the caudate putamen (Ramu *et al*., 2006). Such findings suggest functional reorganization and a broadening of the effects of injury to involve the ipsilesional hemisphere.

### Somatosensory Circuit

In our data, somatosensory cortex (S1, S2) showed broad decreases in glucose uptake. These findings are consistent with prior neuroanatomical studies using both retrograde and anterograde tract tracing which demonstrated projection neurons from the cervical spinal cord to neurons throughout the forelimb area of the sensorimotor cortex and region of secondary somatosensory cortex (S2) (Liang *et al*., 2011; Steward *et al*., 2021). Ghosh et al (Ghosh *et al*., 2009) using retrograde tracers injected after hemisection of the cervical spinal cord, found that corticospinal tract axons of the intact, nonlesioned side sprout across the midline of the spinal cord, with the ipsilesional sensory and motor cortices undergoing important reorganizations. More broadly, hypometabolism was observed in subcortical regions comprising the spinothalamic tract that processes somatosensory input from the limbs and trunk (ventral posterolateral thalamic nucleus), the face (ventroposteromedial thalamus), as well as pain and temperature (ventromedial posterior and mediodorsal nuclei of the thalamus). In addition, hypometabolism was noted in the posterior thalamic nucleus (Po) and the insula that play key roles in multimodal sensory integration and higher-order processing. These regions of the thalamus have been shown to bilaterally project to cervical regions of the spinal cord (Burstein *et al*., 1990). Altered electrophysiological responsiveness have previously been reported post-SCI in the PO, VPL and VPM in rats (Gerke *et al*., 2003; Paulson *et al*., 2005; Hubscher & Johnson, 2006). In principle, the decreased glucose uptake we noted in these thalamic nuclei may be the result of deafferentation. Alternatively hypometabolism in the thalamus may be the result of activation of the ipsilateral reticular nucleus of the thalamus which provides inhibitory, GABAergic input to the dorsal thalamus, including the ventral posterior nuclei (VPL, VPM), lateral dorsal, posterior and ventral lateral nuclei and may have inhibited these structures (Jones, 1975; Jones, 1985). In addition, the reticular nucleus is known to also inhibit neurons of the mesencephalic reticular formation which also showed decreased glucose uptake in our study (Nanobashvili *et al*., 2024).

Of note, both major somatosensory tracts (the spinothalamic/anterolateral tract and the dorsal column-medial lemniscus pathway) from the right side of the spinal cord provide first order structural connections unilaterally to the left VPL. Given that the VPL receives a monosynaptic connection from the contralateral spinal cord, we examined the functional connectivity of the VPL in our dataset (**Fig. 5**). Placement of a region-of-interest in the left (contralateral) VPL demonstrated significant functional correlation in SHAM animals within the extended region of the VPL, both on the left and right side. Though the left and right ventroposterolateral (VPL) nuclei do not share a structural connection through direct commissural fibers, others have in human subjects reported homotopic functional connectivity between bilateral VPL at rest and during motor tasks using fMRI (Mancuso *et al*., 2019; Charyasz *et al*., 2025). This has been proposed to reflect indirect synchronization via corticothalamocortical loops and transcallosal exchanges between homologous somatosensory cortices rather than direct anatomical connections. In SCI animals, in contrast, functional connectivity within the VPL was significantly diminished.

The VPL projects ipsilaterally to the primary somatosensory cortex (S1), with the oral subdivision of the VPL also sending thalamocortical axons to the ipsilateral primary motor cortex (M1). SCI animals in contrast to SHAM animals demonstrated a significant loss of functional connectivity to bilateral somatosensory cortex (S1, S1BF, S1FL, S1HL, S1ULp) and motor cortex (M1)(second order functional connections: Spinal cord→VPL→S1, M1), which themselves are known to have reciprocal monosynaptic structural connections to the VPL (Horne & Tracey, 1979; Shook *et al*., 1991; Clasca, 2023; Rubio-Teves *et al*., 2024). This is consistent with prior work demonstrating using fMRI that functional connectivity between VPL and S1 is decreased 1 week after unilateral, cervical SCI, suggesting a disruption of second order synaptic connections (Seminowicz *et al*., 2012).

### Motor circuit

Decreases in glucose uptake were noted in regions of the motor circuit, including primary motor cortex, striatum (dorsal, central, ventral), globus pallidus, subthalamic nucleus and the substantia nigra. Overall, the hypometabolism in motor cortex was less broadly represented than that in somatosensory cortex. Hypometabolism in the striatum was broadly seen in ventral regions, but in dorsal regions was significant mostly posteriorly. Prior work using anterograde and retrograde molecular tracers, as well as viral tract tracers has confirmed projections of the cervical spinal cord to motor cortex, striatum, globus pallidus and the subthalamic nucleus (Newman *et al*., 1996; Liang *et al*., 2011; Kamiyama *et al*., 2015; Poinsatte *et al*., 2024).

In contrast, a small significant increased glucose uptake was noted bilaterally in posterior secondary motor cortex, and in the contralateral ventrolateral (VL, motor) thalamus. Increases in activation of the contralateral VL assessed by perfusion changes have previously been reported by Morrow et al. after T12 to L3 spinal lesions (Morrow *et al*., 2000; Paulson *et al*., 2005). Noteworthy, was that the cerebellum primarily showed bilateral increases in regional glucose uptake, including the lateral lobes (cruz1), lobules 4 and 5 of the vermis, the interposed and lateral deep cerebellar nuclei, as well as the middle cerebellar peduncle. The dorsal spinocerebellar tract projects mainly ipsilateral to the cerebellum, but a contralateral projection is also present (Matsushita & Xiong, 1997). Prior work in clinical SCI patients using [^18^F]-fluoro-2-deoxyglucose positron emission tomography (^18^F-FDG PET) has also confirmed increased glucose uptake in the cerebellum (Curt *et al*., 2002b; Chopra *et al*., 2022), with changes also reported in cerebellar synaptic markers (Visavadiya & Springer, 2016).

### Limbic circuit

Different from somatosensory and motor regions, limbic areas after hemicontusive SCI prominently showed widespread increases in glucose uptake alongside select areas of hypometabolism. Increases were noted in the amygdala (basolateral n., medial n., cortical amygdala), nucleus accumbens (core, shell), lateral septum, hypothalamus (posterior), and hippocampus (CA1-3, DG). The functional changes in limbic regions are consistent with earlier studies showing that labelling of the amygdala and nucleus accumbens neurons with fluorogold showed retrograde uptake of the tracer throughout the spinal cord, in particular the upper cervical and lumbar segments (Burstein & Giesler, 1989; Burstein & Potrebic, 1993). Anterograde and retrograde transport studies from the upper cervical spinal cord and the cervical enlargement have also demonstrated direct spinal projections to the amygdala, septum and hypothalamus (Newman *et al*., 1996; Liang *et al*., 2011). Increased activation of the nucleus accumbens has been previously described as an important component of recovery after hemicontusion of the cervical spinal cord (Sawada & Nishimura, 2021).

In addition, our study showed decreases in glucose uptake in regions of the thalamus associated with processing of emotional information, including anterior midline structures (anterior, paraventricular), mediodorsal, habenular, as well as lateral dorsal nuclei. Prior evidence for structural or functional changes in limbic-related nuclei (mediodorsal, anterior, midline/intralaminar, lateral dorsal) after SCI is sparse. Metabolic imaging hints at widespread thalamic alterations, but targeted studies on emotional/limbic thalamic nuclei post-SCI remain an open area for research.

### Clinical correlates

The occurrence of SCI sets into motion a set of neuronal, glial, neuroplastic, and inflammatory responses that take months to evolve. During this time, factors including the dying back of neurons, proliferation and activation of glial cells, compensatory and functional reorganization of neural circuits, shape regional cerebral metabolic needs. Changes in functional reorganization can occur as early as hours to days after injury (Sydekum *et al*., 2014; Yagüe *et al*., 2014). During the transition from acute to chronic SCI, neural circuits may undergo changes in functional reorganization that differ across brain regions. For example, Jurkiewicz et al. reported that when SCI patients were imaged longitudinally during performance of a simple motor task shortly after injury and continuing over the course of 1 year, patients compared to healthy controls demonstrated increasing M1 activation, whereas associated sensorimotor areas showed a progressive decrease in activation (Jurkiewicz *et al*., 2007).

In humans, most neuroimaging studies have focused on the chronic phase of SCI. Here, similar to our results, studies have reported a diffusion of functional activation (cerebral perfusion-based) to include the unaffected limb representation (Curt *et al*., 2002a), as well as activation outside of the expected pattern of activation evoked by the particular motor task (Bruehlmeier *et al*., 1998). Increased activation of the cerebellum has also been reported in SCI patients during a motor challenge (Bruehlmeier *et al*., 1998; Curt *et al*., 2002b). ^18^F-FDG-PET brain imaging in SCI patients has shown variable results. Some have reported reduced metabolic uptake in sensory cortical areas, anterior cingulate gyrus, and hippocampus (Chopra *et al*., 2022), others have reported such reductions in midbrain, the cerebellum and the temporal cortex (Roelcke *et al*., 1997). Raised metabolism has been reported in supplementary motor areas (SMA), inferior frontal gyrus, some areas in the parietal and temporal lobe, putamen and cerebellum (Chopra *et al*., 2022), or in SMA, anterior cingulate cortex and the putamen (Roelcke *et al*., 1997). Differences may relate to patient selection (paraplegia, tetraplegia), time since injury, as well as whether imaging was performed at rest or during a motor task. Functional brain reorganization after SCI has been implicated in a number of its debilitating sequelae. Reorganization of sensory circuits is proposed to contribute to hyperalgesia, while changes in limbic structures may contribute to depression, a frequent sequelae of SCI. Improved understanding of the interaction of the brain-spinal cord axis, as well as in structure-function relations may have important clinical implications for the evaluation and treatment of patients with SCI.

### Limitations

Our study focused on the subacute phase (1 week) following SCI. We did not study longitudinal changes in neuroplasticity in remote brain networks. Understanding of the temporal course of brain changes may help in the future to define optimal timepoints for neurorehabilitative interventions. We did not examine brain functional activation during a motor or sensory challenge. Such challenges have the potential to unmask latent deficits not appreciated in the resting state. This may help explain the absence of metabolic changes in some regions that previously have been reported to demonstrate structural connections via tract tracing (Liang *et al*., 2011). Our findings of remote changes in cerebral glucose uptake after SCI does not directly address the mechanism of change, and whether this is induced by neuroinflammation, changes in glial cell activation (Wu *et al*., 2014a; Wu *et al*., 2014b; von Leden *et al*., 2016; Wu *et al*., 2016), mitochondrial function (Sullivan *et al*., 2007; Wu *et al*., 2020; Seira *et al*., 2024), decreases in neuronal viability or a downregulation of glycolytic or aerobic pathways, or dysregulation of neural activity. Neither did we examine sex differences (McFarlane *et al*., 2020; Lee *et al*., 2023; Mojarad *et al*., 2025; Pacheco *et al*., 2025) or strain differences (Paulson *et al*., 2005) which are beginning to be reported. Future work could consider direct examination of behavioral correlates, including sensory hyperalgesia, as well as allodynia-like hypersensitivity and their relationship to functional brain changes.

## Data statement

Data access requests should be sent to the corresponding author and will be granted with the signing of a data sharing agreement.

